# Pupation site preference selection in *Drosophila jambulina*

**DOI:** 10.1101/333088

## Abstract

Larvae of *Drosophila jambulina* belonging to montium subgroup were tested for pupation site preference in relation to temperature. At higher temperature (30 °C), larvae preferred to pupate on food whereas at lower temperature (21 °C) pupation occurred on the cotton. Genetic basis of larval pupation behavior was studied by conducting reciprocal crosses for 30 generations on food-selected and on cotton-selected larvae. Results from genetic analysis between food-selected and cotton-selected strains suggested a single gene responsible for the pupation site preference, with F1 progeny pupated on cotton and F2 (F1×;F1) larvae pupated on both food as well on cotton. Although we found no change in morphological traits in food *vs.* cotton selected population, significantly different growth rate (body weight) between the two strains was observed. These results suggest that pupation site preferences can affect life-history traits in *D. jambulina.*

## 1. Introduction

A behavioral phenotype is influenced by both genotype and environment (Sokal et al., 1960). Difference in pupation site choice can reflect larval difference in the method of finding food as well as difference in niche breadth (Jaenike, 1985). Since the larvae of *Drosophila* inhabit in a challenging environment surrounded by various parasites, predators, abiotic stresses such as excessive heat and cold, prior to pupation larvae initiate wandering behavior to find the thermally favorable locations for pupation (Dillon et al., 2009). For example, it has been observed that *Drosophila* tends to pupate close to the food when temperatures are high whereas at lower temperatures pupation occurs high on the vial sides (Schnebel and Grossfield, 1992). However, these pupation height differences may not depend on temperature *per se*, rather on correlates of temperature, such as the moisture content of the food and the humidity of the air (Sokal et al., 1960).

Previous studies have revealed that pupation site preference (PSP) in various species of *Drosophila* is affected by both biotic (sex, density, locomotory path length, developmental time and digging behavior) and abiotic (temperature, humidity, moisture, light and dark and pH) factors (Sokal et al., 1960; de Souza et al., 1970; Fogleman and Markow, 1982; Pandey and Singh, 1993; Hodge et al., 1996; Hodge and Caslaw, 1997). Additionally, genetic factors control the pupation behavior in various species of *Drosophila* (de Souza et al., 1970; Bauer and Sokolowski, 1985, 1989; Garcia-Florez et al., 1989). For example, Joshi and Muller (1993) reported that pupation height is a polygenic trait that responds effectively to bidirectional selection. Furthermore, it has been found that preference of larvae to pupate on food or at the bottom of culture bottles is because of difference in a single gene (de Souza et al., 1970). Our primary aim in this study is to establish the genetic basis of laboratory selected strain with food pupation site preference and cotton site preference in *D. jambulina*. Further we intended to analyze effects of direct selection on traits related to morphology and life-history after every five generations. Conclusions of the present study are based on i) maintenance of selection experiment in laboratory for 30 generations, ii) measurement of genetic variation in the traits after every five generations.

## 2. Materials and methods

### 2.1. Laboratory selection

#### 2.1.1 Habitat selection

Samples of wild *D. jambulina* (*n* = 100-120) were collected from three geographically distinct sites of the Indian subcontinent (Table 5). Samplings were done in months of February-March using net sweeping and bait trap method from fruit markets, godowns and nurseries from each of the localities. Based on the average temperature of the sampling localities, cultures were maintained at 21 °C. To maintain the low density (30-40 eggs per vial) egg laying period was restricted for 6 to 8 hours. Climatic data of sampling localities were taken from the website of Indian Institute of Tropical Meteorology (IITM; www.tropmet.res.in).

#### 2.1.2 Food pupation and cotton pupation site

To carry out the selection experiment, four different stocks were maintained each with sufficient variation for pupation site preference. For each stock, 50 pairs of arbitrarily selected flies were isolated in separate wide-mouth bottles (300 mL capacity). Additionally, a control population was maintained and was not subjected to any selection regime. In the selection experiment, 100 males and 100 females derived from food pupae as well as from cotton pupae from each generation were selected to initiate the next generation. Selection method for each of the mass-bred stocks was followed independently. This selection procedure was followed independently for each of the mass-bred stocks. Selection experiment was carried out for 30 generations which resulted in food-selected strains (L1, L2, L3, and L4) and cotton-selected strains (H1, H2, H3, and H4). Replicates of each of the selected strains (L1, L2, L3, and L4 and H1, H2, H3, and H4) were kept under relaxed selection after the 30th generation i.e., from the 31st through the 35th generation. These replicate strains were maintained with the same number of flies without any further selection regime. The response of the selection was measured after every fifth generation.

### 2.2 Genetic crosses

To determine the genetic basis of pupation site preference and allelic dominance, a set of eight crosses between laboratory-selected true-breeding food strain (L) and cotton strain (H) differing in pupation behavior was performed. Each of the eight crosses was made using virgin1H♀ and 1L ♂ to obtain F1 progeny.

To understand the pattern of inheritance, reciprocal crosses (virgin 1L ♀ and 1H ♂) were also set in equal numbers. The F1 progeny were then checked for pupation site preference. To obtain F2 generation, 50 ♂ and 50 ♀ (five sets of 10 pairs each) were randomly selected from L1 progeny. After pupation, preference site of larvae were examined and number of pupae on each site (food *vs*. cotton) were counted.

### 2.3 Correlated responses to selection: analysis of morphological and life-history traits

The correlated selection changes in morphology, wing length, thorax length and wing width were examined. Additionally, changes in life-history traits in food selected and cotton selected replicate lines of *D. jambulina* were analyzed.

#### 2.3.1 Measurement of body size

Measurement of wing length, wing width and thorax length was done under Olympus stereozoom microscope SZ-11, Japan (www.olympus.com) fitted with a micrometer. Wing length was measured from the thorax articulation to the tip of third longitudinal vein whereas wing width was measured at the middle of the wing where the width of the wing was the maximum. Length of thorax was measured from anterior margin up to the tip of postscutellum.

### 2.4 Statistical analysis

Chi square (χ^2^) test was used to determine the goodness of fit between observed and expected number of larvae pupated on food and on cotton. The data was subjected to *t*-test to analyze the difference between food selected and cotton selected strains for various morphometrical and life-history traits. For each population, frequency of pupation (in %) on food and on cotton for cotton – selected and food-selected strains and their reciprocal F1 are presented as mean values.

## Results

### 3.1 Laboratory selection of food selected pupae and cotton selected strains

For selection regime, populations having maximum viability in pupation site preference (PSP) were used. Selection for PSP in *D. jambulina* for 30 generations resulted in a two different phenotype (larvae pupate on food) in food selected strains and (larvae pupate on cotton) in cotton-selected strains (Figure 1).

**Figure 1.**
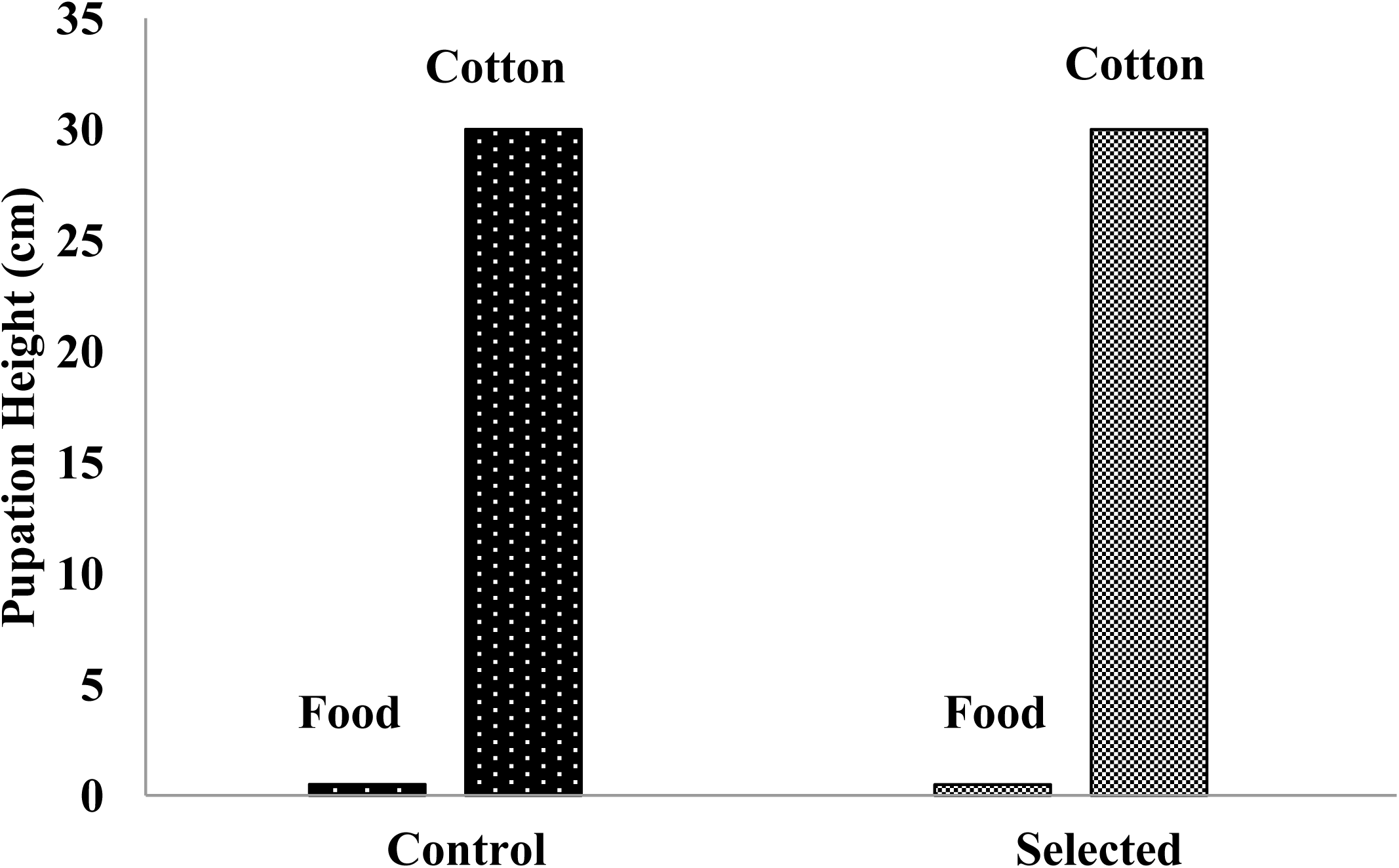
Graph showing no change in pupation height after selection.

**Figure 2.**
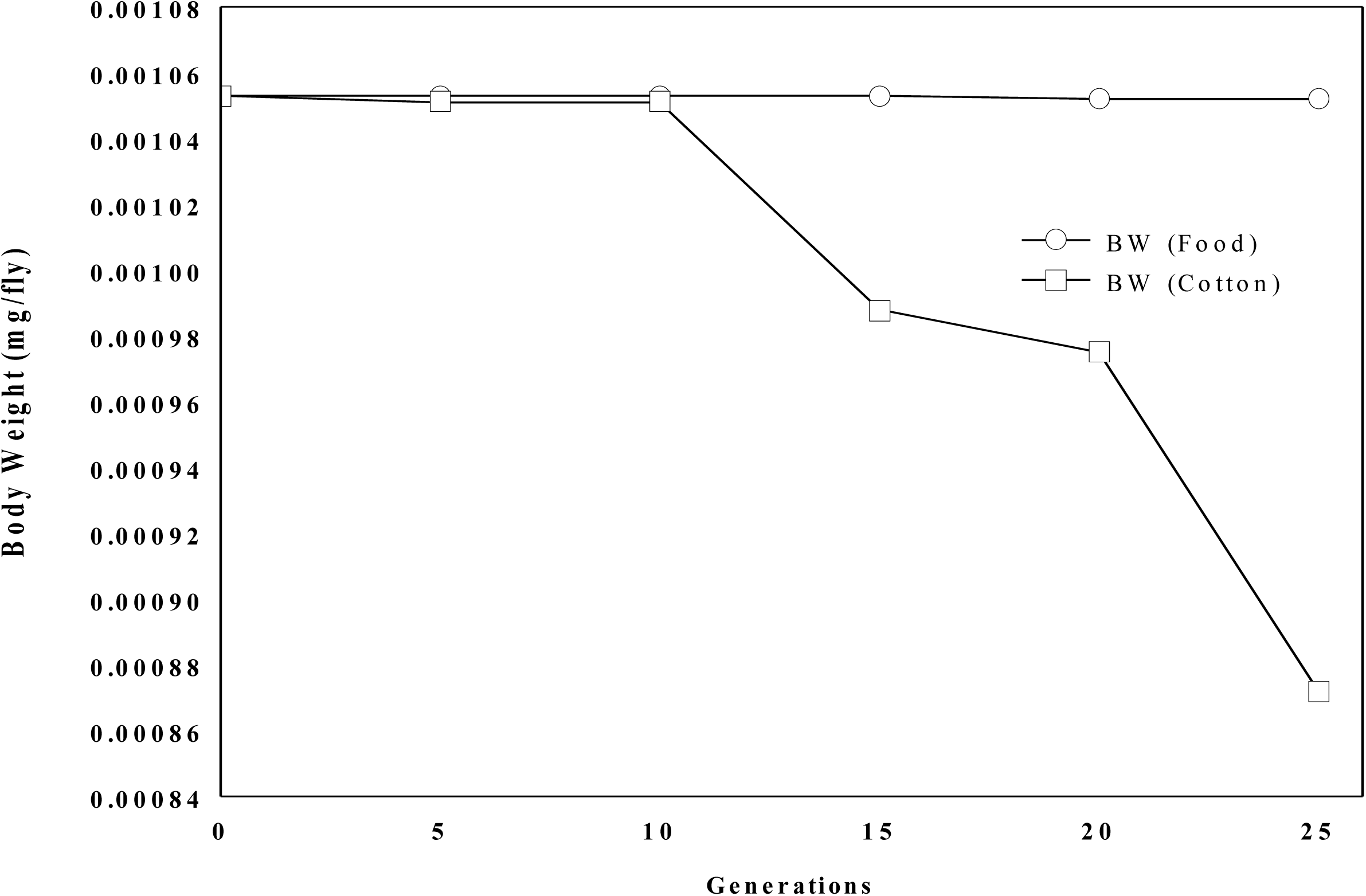
Graph showing change in body weight in selected strain with successive generations.

**Figure 3.**
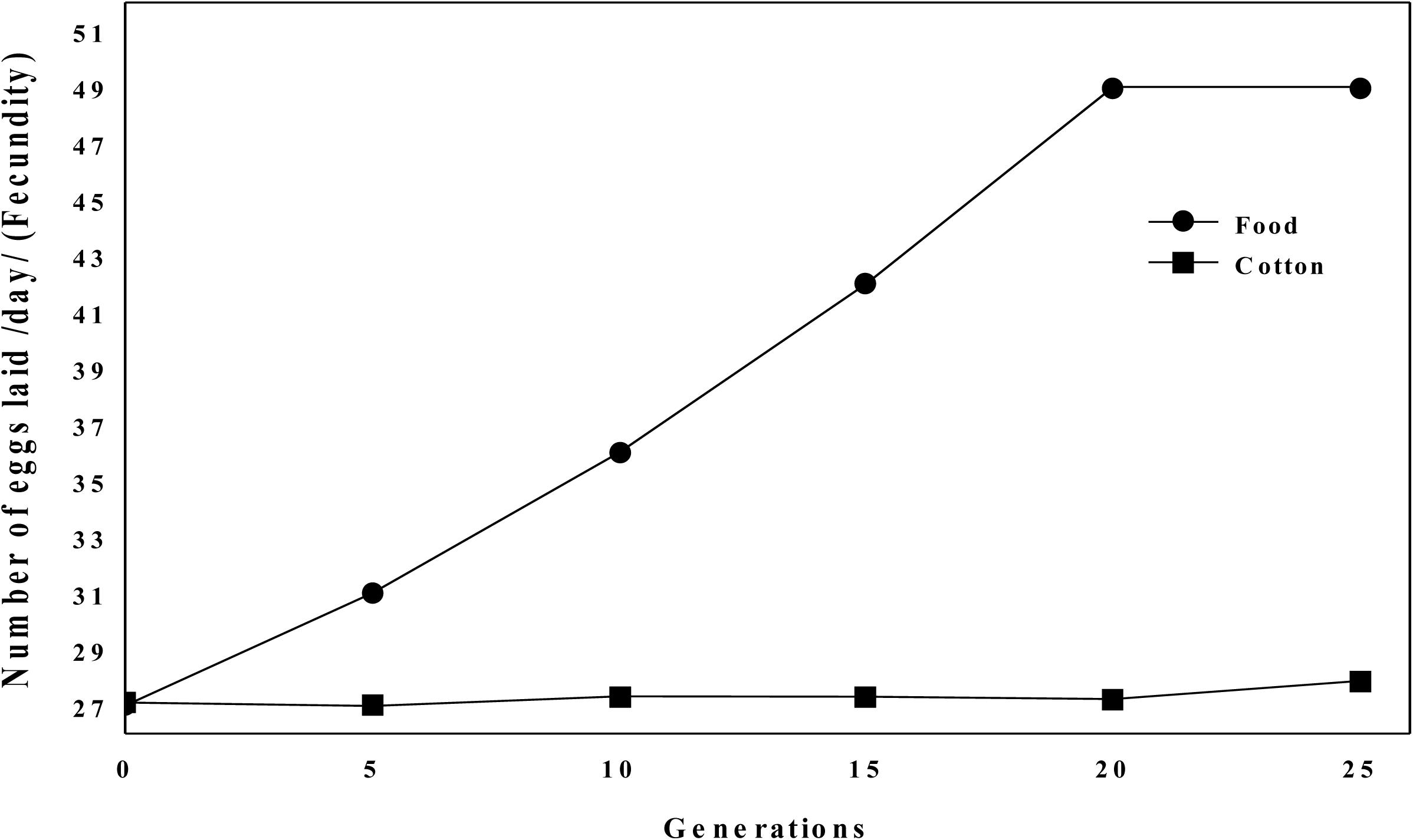
Correlated changes in fecundity are shown after every five generations of laboratory selection in food selected and cotton selected strains. Selection was made on the pupae.

### 3.2 Genetic basis of pupation site preference

True breeding strains were used for cotton-selected (HH) and food-selected (LL) strains to examine the genetic basis as well as allelic dominance for pupation site preference. Reciprocal crosses for F1 progeny always pupated on cotton (Table1). In reciprocal F2 crosses (F1×;F1), larvae pupated on both food as well on cotton and phenotypic ratios of F strain and L strain pupae followed Mendelian F2 ratio (Table1). These results showed Mendelian mode of inheritance of a single gene with two alleles.

**Table 1.**
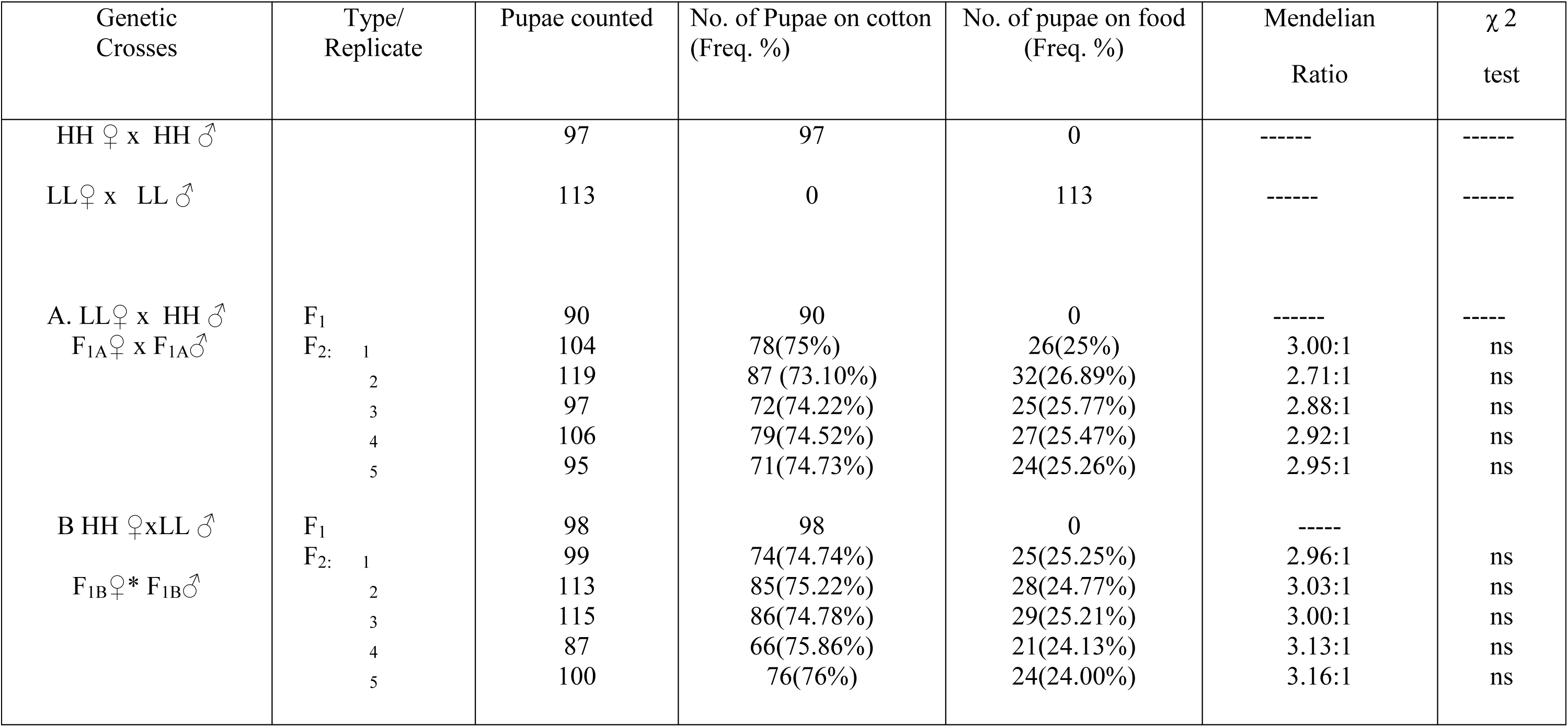
Data of pupation site preference in F1 and F2 progeny (pupae) of F1 and F2 genetic crosses between laboratory-selected true breeding food-selected (LL) and cotton-selected (HH) strains of the fruit fly *Drosophila jambulina* at 21 °C growth temperature.

| Genetic Crosses | Type/<br>Replicate | Pupae counted | No. of Pupae on cotton<br>(Freq. %) | No. of pupae on food<br>(Freq. %) | Mendelian<br>Ratio | $\chi^2$<br>test |
| --- | --- | --- | --- | --- | --- | --- |
| HH ♀ x HH ♂ |  | 97 | 97 | 0 | ----- | ----- |
| LL ♀ x LL ♂ |  | 113 | 0 | 113 | ----- | ----- |
| A. LL ♀ x HH ♂ | F<sub>1</sub> | 90 | 90 | 0 | ----- | ----- |
| F<sub>1A</sub> ♀ x F<sub>1A</sub> ♂ | F<sub>2</sub>: 1 | 104 | 78(75%) | 26(25%) | 3.00:1 | ns |
|  | 2 | 119 | 87 (73.10%) | 32(26.89%) | 2.71:1 | ns |
|  | 3 | 97 | 72(74.22%) | 25(25.77%) | 2.88:1 | ns |
|  | 4 | 106 | 79(74.52%) | 27(25.47%) | 2.92:1 | ns |
|  | 5 | 95 | 71(74.73%) | 24(25.26%) | 2.95:1 | ns |
| B HH ♀xLL ♂ | F<sub>1</sub> | 98 | 98 | 0 | ----- |  |
| F<sub>1B</sub> ♀* F<sub>1B</sub> ♂ | F<sub>2</sub>: 1 | 99 | 74(74.74%) | 25(25.25%) | 2.96:1 | ns |
|  | 2 | 113 | 85(75.22%) | 28(24.77%) | 3.03:1 | ns |
|  | 3 | 115 | 86(74.78%) | 29(25.21%) | 3.00:1 | ns |
|  | 4 | 87 | 66(75.86%) | 21(24.13%) | 3.13:1 | ns |
|  | 5 | 100 | 76(76%) | 24(24.00%) | 3.16:1 | ns |

### 3.3 Changes in dominance of pupation site preference

Pupation preference of selected strains were not affected by the temperatures whereas pupation site preference of F1 (HL and LH) and F2 were strongly affected by the temperature (Table 2). Larvae of F1 (HL and LH) and F2 favor to pupate on cotton at 21 °C and 80% relative humidity (Rh) whereas at higher temperature (30°C and 80% Rh) larvae of F1 (HL and LH) prefer to pupate on food. The change in temperature caused shift in Mendelian Ratio (3 cotton selected: 1 food selected) at 21 to (3 food selected: 1 cotton selected) at 30°C.

**Table 2.**
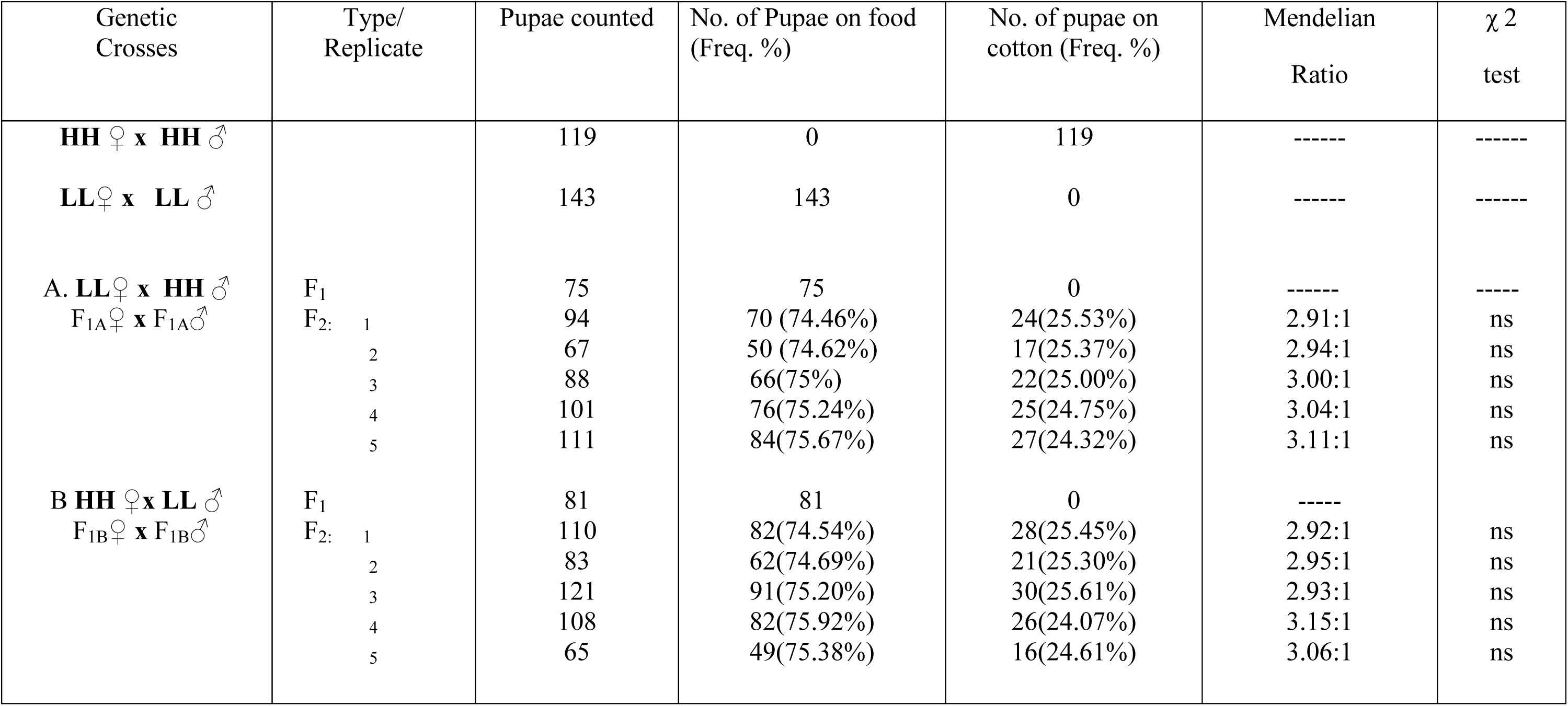
Data of pupation site preference in F1 and F2 progeny (pupae) of F1 and F2 genetic crosses between laboratory-selected true breeding food-selected (LL) and cotton selected (HH) strains of the fruit fly *Drosophila jambulina* at 30 °C growth temperature.

### 3.4 Effect of selection on morphology, rate of development, life-history traits

Morphological traits (wing length, wing width and thorax length), did not respond to selection in *Drosophila jambulina* (Table 3). Similarly, overcrowding experiment with 100 pairs of flies per bottle did not show any change in percent frequency of pupation site preference. The degrees to which larvae showed their pupation site preference were 100% in selected directions.

**Table 3.** Mean (± S.D.) values for various morphometrical traits and life history trait in control and selected strain of *D. jambulina*. For significance level between control and selected strain student’s *t-* test values were used.

| Trait | *D. jambulina* |  |  |  |
| --- | --- | --- | --- | --- |
|  | Control | Food | Cotton | *t*-test |
| Wing length (mm) | 2.05 ± 0.05 | 2.05± 0.04 | 2.05±0.06 | ns |
| Thorax length (mm) | 1.00± 0.04 | 1.00± 0.05 | 1.00±0.03 | ns |
| Wing width (mm) | 0.77± 0.02 | 0.77± 0.03 | 0.77±0.02 | ns |
| Body weight (mg/fly) | 0.001053 | 0.001153 | 0.000872 | ** |
| Fecundity /day | 27 | 38 | 25 | ** |
| DOD (Days) | 16 | 13 | 19 | *** |

To study the effect of selection on body weight, ten adult files (7 days after eclosion) per selected strain of each sex were weighed. Significant variation (*P* < 0.001) in the body weights of the selected strains was observed. The food-selected flies were always heavier than cotton-selected flies of the same sex raised in the same environment (Table 4). Rate of development was also affected by the selection; food-selected flies took 14 days from the laying of the eggs to eclosion while it was 16 days in cotton selected flies.

**Table 4.**
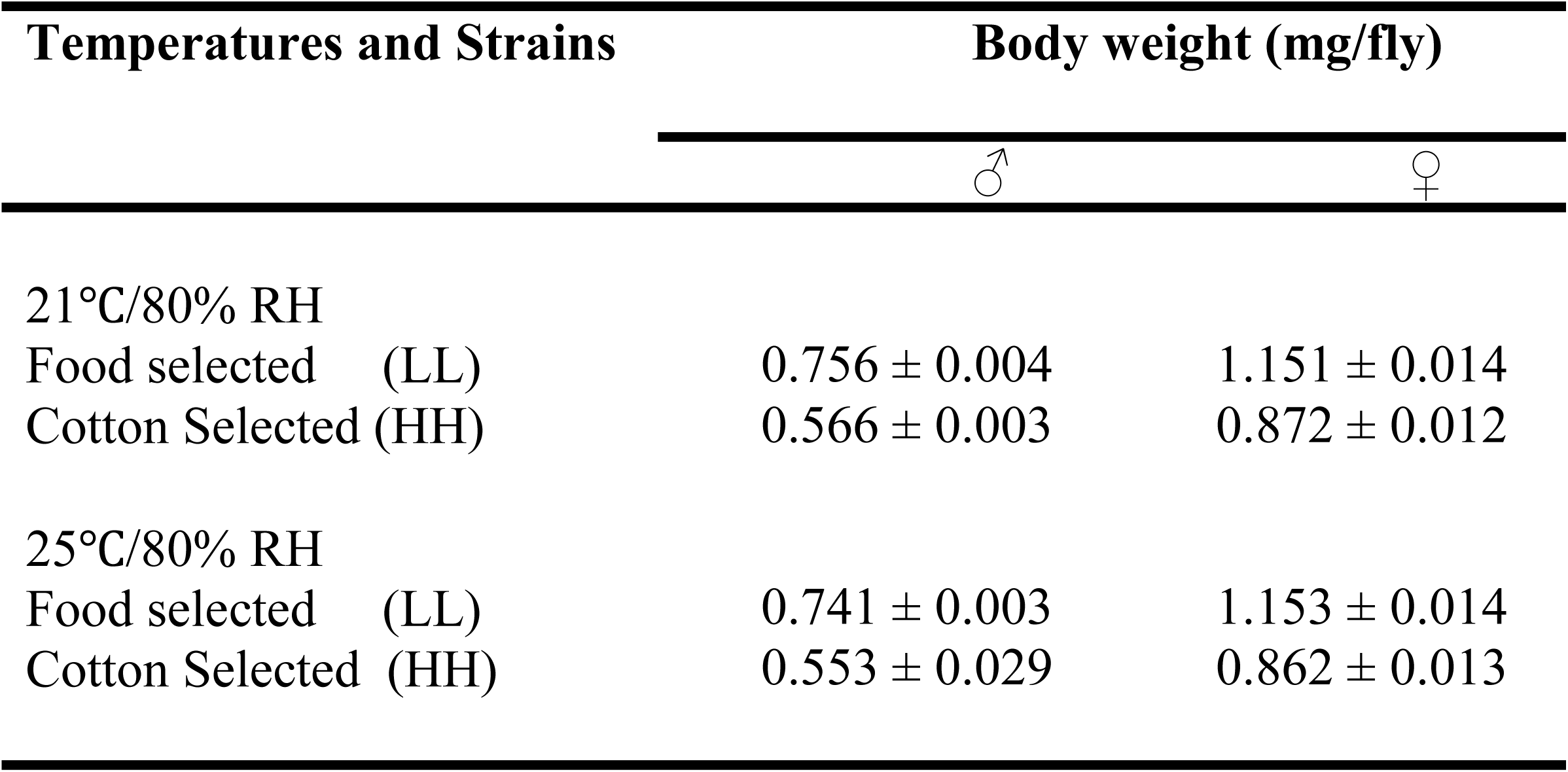
Mean weights (mg/per fly) ± S.E of each sex from each temperatures.

### 3.5 Geographical difference in pupation site preference

Larvae derived from natural populations showed variation in pupation site preference corresponding to geographical populations. Data of pupation site preference in three geographically distinct populations of *D. jambulina* are shown in table 4. The F2 progeny (pupae) of reciprocal genetic crosses of food-selected and cotton-selected strains resulted in pupation site preference differences in two classes (Table 1, 2). Therefore, results of F2 crosses were checked in the wild-caught samples (Table 5). Contrasting patterns of pupation site preference in northern versus southern populations of *D. jambulina* were found. Frequency of pupae on food was three times higher than on cotton in northern populations (Rohtak and Kalka) whereas in southern population (Trivandrum) a reverse trend was observed with three times more frequency of pupae on cotton than on food.

**Table 5.** Data on wild-caught females (pupae) for variation in pupation site preference in the fruit fly *Drosophila jambulina* populations latitudinal sites on the Indian subcontinent.

| Geographical site (latitude) | No. of pupae | Pupae on food (freq. %) | Pupae on cotton (freq. %) | Mendelian Ratio | $\chi^2$ test |
| --- | --- | --- | --- | --- | --- |
| **Trivandrum (8.29°N)** | 73 | 19 (26.02%) | 54(73.97%) | 2.84:1 | ns |
| **Rohtak (28.35)** | 168 | 130(77.38%) | 38(22.61%) | 3.42:1 | ns |
| **Kalka (30.50°N)** | 64 | 47(73.43%) | 17(26.56%) | 2.76:1 | ns |

## 4. Discussion

In this study, we have examined the effect of growth temperatures and its genetic basis on laboratory-selected strains of *Drosophila jambulina* for pupation site preference (PSP). Results showed a considerable variation in pupation site preference in natural populations corresponding to temperature variation. Previous studies have shown that temperatures can affect larval behavior which may cause significant variation in PSP (Rodriguez and Sokolowski, 1987; Schnebel and Grossfield, 1986, 1992; Vandal and Shivanna, 2007). Our results are consistent with these studies, as we found that frequency of pupae pupating on food and on cotton varied with temperatures. At higher temperature (30 °C), larvae pupated on food whereas at lower temperature (21 °C) pupation occurred on the cotton used to plug the mouth of the culture bottles. The variation observed in the PSP may be attributed to amount of synthesis of glue protein. Higher temperature (30 °C) acts as a inducer for the synthesis of large quantities of glue protein and helps the pupae to stick to the food for the pupation while at lower temperature (21 °C) decrease in quantity of synthesized glue protein allow larvae to move along with sides of container for pupation on cotton.

The frequency of F2 progeny (pupae) of genetic crosses between food-selected and cotton-selected strains in our study was close to Mendelian ratio suggesting that inheritance of PSP fit a single-gene locus with two alleles. However, inheritance of pupation behavior may differ between species. For example, inheritance of PSP in *Drosophila melanogaster* was found to have additive polygenic model of inheritance (Bauer and Sokolowski, 1988). The disparity in pupation behavior in different species could be due to different adaptive values in their gene pools.

Although limited number of traits can be studied through artificial selection experiments, they can directly reveal patterns of responses and co-responses of selected traits (Ramniwas et al., 2013). For *D. jambulina*, rapid response to direct selection of pupation site confirms the presence of genetic variation for the selected trait. Furthermore, results suggest pleiotropic effects of some of the genes that control the variation in pupation site preference. As in our experiment we found association of pupation behavior with fitness of the flies.

Our results show that having a strong preference for either site (food or cotton) affect individual’s fitness. Flies eclosed from cotton-selected pupae showed decrease in body weight while body weight was observed in food-selected pupae. The difference in weight could be attributed to the fact that food-selected pupae get all the food for themselves without any competition as cotton larvae leave the food two days before pupation. Morphological traits (wing length, wing width and thorax length), did not respond to selection in *Drosophila jambulina.* However, growth temperature caused change in allelic dominance. This could be due to different adaptive values of alleles in different environment.

## Conclusions

In the study, a rapid response to bidirectional selection for food-selected and cotton-selected strains in *D. jambulina* was observed. Genetic crosses between laboratory-selected food and cotton strains showed 3:1 Mendelian ratio in both the crosses (F2). Our data fit a single-gene locus model for pupation site preference in *D. jambulina*. Selected strains lack effect of temperature, but in F1 and F2 progeny (pupae) effect of temperature was clearly visible. Selected strains of *D. jambulina* developed an adaptive behavioral polymorphism. Additionally, we found that difference in pupation site preference affect life-history traits in *D. jambulina*.

## Acknowledgements

The first author is grateful to the Department of Science and Technology (DST), New Delhi, for the financial support through the DST-INSA/INSPIRE faculty fellow project (IFA-11LSBM-08). Thanks are also due to Professor Sanjeet Singh, Associate Dean Research, Chandigarh University for the help and support to carry out the research.

